# Rapid trait adaptation has contrasting effects on different dimensions of stability under environmental change

**DOI:** 10.64898/2026.10.02.756195

**Authors:** Mark Holmes, Masato Yamamichi, Frederik de Laender

## Abstract

Environmental change alters species growth rates and, through species interactions, can produce complex effects on community functioning and composition. Because functional traits mediate both environmental responses and competition, rapid trait adaptation may modify these effects. We used a trait-based model of competitive communities to test how adaptation affects the mean and diversity of species growth responses to environmental change, and how this influences functional and compositional resistance and resilience. Adaptation altered response diversity more strongly, by changing the balance between environmental optimisation and competition-driven trait divergence. As a result, adaptation strongly increased compositional stability, but had less clear effects on abundance. Adaptation also generated a trade-off between resistance and resilience: faster adaptation improved resistance but reduced resilience as communities were displaced less by the environmental change and thus recovered less. Overall, adaptation can be stabilising, but faster adaptation does not necessarily produce greater stability across all stability components.

## Introduction

Anthropogenic activities alter environmental conditions, for instance by increasing temperature or pollutant concentrations. These changes cause direct effects on species growth rates which, propagated through species interactions, can cascade to unexpected net effects on ecosystem functioning (e.g. total community abundance) and community composition (Arnoldi, Loreau, and Haegeman 2019; Loreau et al. 2001; Valladares et al. 2015). For instance, even modest reductions in precipitation can alter competitive dynamics between plant species, reshaping coexistence outcomes beyond what direct growth-rate effects alone would predict (Van Dyke, Levine, and Kraft 2022). Understanding the mechanisms leading to such net effects is essential to predict changes in ecosystem services such as biomass production.

The direct effect of environmental change on species growth rates can be summarised as both the mean response and the variation across responses (De Laender et al. 2023; Laloux and De Laender 2026; Ross et al. 2023), the latter often referred to as “response diversity”. In competitive communities, species interactions are typically expected to lead to greater loss of total abundance when the mean response is large, while greater variation in species responses would result in more pronounced compositional change (Holmes, Spaak, and De Laender 2021; Laloux and De Laender 2026; Pitzer et al. 2026).

Functional traits can help predict community-level effects of environmental change when they mediate both species’ direct responses and the interaction strengths which translate those responses into community-level outcomes (Suding et al. 2008). One example of such a functional trait is body size, which often influences responses to changing environmental conditions (Brown et al. 2004; Echeveste, Agustí, and Dachs 2010; Gillooly et al. 2001; Kremer, Thomas, and Litchman 2017; Mensens et al. 2017) and dictates the strength of competition (Edwards, Litchman, and Klausmeier 2013; Gallego, Venail, and Ibelings 2019; Sebastián-González et al. 2017; Van Donk, Ianora, and Vos 2011). Integrating both species responses and species interactions within a trait-based model is therefore a natural approach to studying net effects of environmental change (Barabás and D’Andrea 2016; Barabás et al. 2022).

Functional traits are not static, but can change with time through adaptive processes, including genetic evolution and phenotypic plasticity (Fusco and Minelli 2010; Kopp and Matuszewski 2014). Such trait change may be rapid enough to proceed at a rate comparable to ecological processes (Ellner, Geber, and Hairston 2011; Hairston et al. 2005; Yamamichi and Letten 2021) and therefore modify both direct effects of the environment and species interactions on the same time scale. If and how rapid adaptation affects species growth responses to environmental change, and how this influences stability, is at present unknown, since this question has mostly been studied in one- and two-species communities (Catullo et al. 2019; Morita and Yamamichi 2023). In this paper, we therefore examine how trait adaptation affects species responses, community functioning (abundance), and community composition, in competitive communities exposed to environmental change. We consider three kinds of empirically measurable ecological stability metrics (Bowler et al. 2020; Clark et al. 2021; Donohue et al. 2016; Halpern et al. 2008; Hillebrand et al. 2018; Jarillo, Cao-García, and De Laender 2022; Radchuk et al. 2019): resistance, resilience, and recovery. Resistance quantifies how unaffected a community is by environmental change, while resilience quantifies the rate at

which it returns towards its original state (Hillebrand et al. 2018; Pimm 1984; Van Meerbeek, Jucker, and Svenning 2021). Because we consider environmental change that is sustained (i.e,. a press perturbation) rather than temporary (i.e,. a pulse perturbation), the community need not return to its pre-disturbance state. Here, we interpret resilience as the transient tendency to move back *toward* the pre-disturbance state relative to the initial displacement. Finally, recovery quantifies the absolute difference between the final and pre-disturbance states (Van Meerbeek, Jucker, and Svenning 2021).

We ask three questions. First, how does adaptation affect the mean and variation of species growth responses? Second, how does the rate of trait adaptation affect functional and compositional resistance? Third, does adaptation permit functional or compositional resilience and recovery in a scenario of sustained environmental change? We quantify these stability metrics both in terms of functioning (total community population) and composition (similarity to unstressed communities).

We find that the effects of adaptation on response mean and variation depend on the relative importance of competition and environmental change on growth. Adaptation strongly improves compositional resistance, while having only limited ability to mitigate the effects of environmental change on growth and, subsequently, functioning. In scenarios of sustained environmental change, adaptation indeed allowed some degree of compositional recovery, however this is inversely related to the rate of adaptation, as more-adaptive communities are able to adapt more smoothly towards their final state.

## Methods

### Community model

We use a generalised Lotka-Volterra model incorporating trait-dependent growth and trait adaptation. The rate of change of the abundance of each species *i, N*_*i*_, within the community of *n* species depends on both the amount of environmental change, *E*, and the value of the species’ quantitative trait, *X*_*i*_, and that of its competitor, *X*_*j*_:

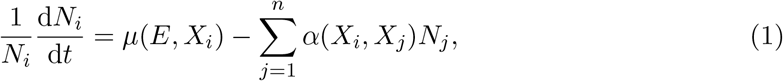

where *µ*(*E, X*_*i*_) is the intrinsic growth rate and *α*(*X*_*i*_, *X*_*j*_) is the trait-based interaction coefficient between species *i* and *j*.

Environmental change is bounded 0 *≤ E < ∞*, where *E* = 0 indicates completely pristine conditions and *E* = *∞* represents completely degraded conditions. The intrinsic growth rate depends on both *E* and *X*_*i*_:

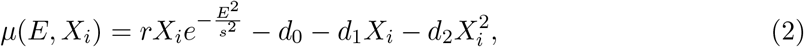

where *r* is the maximal intrinsic growth rate, and *d*_0_, *d*_1_, and *d*_2_ are constant, linear, and quadratic mortality terms with respect to the trait value (see Figure S1). Environmental dependence of the intrinsic growth rate is encoded by multiplying *r* by a Gaussian function with a standard deviation of *s*. A higher *s* means that a larger amount of environmental change is required to reduce the growth rate to some set amount.

Finally, the strength of the competitive interactions scales inversely with the distance between trait values via a Gaussian function (Barabás and D’Andrea 2016; Morita and Yamamichi 2023):

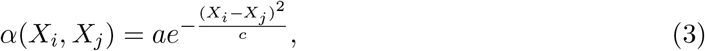

with a variance of *c*; *a* is a scaling factor. This function is such that *α*(*X*_*i*_, *X*_*i*_) = *a* i.e. self-limitation remains constant over time and across species and that max *α*(*X*_*i*_, *X*_*j*_) = *a*. The maximum abundance that species *i* can attain in a given environment (i.e. its carrying capacity, *K*_*i*_(*E, X*_*i*_)) is defined as:

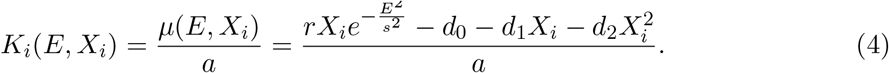

The trait dynamics are based on quantitative genetic models (Abrams 2001; Cortez and Patel 2017; Ellner, Geber, and Hairston 2011; Yamamichi and Letten 2021). The rate of trait change depends on the partial derivative of the per-capita growth rate with respect to the trait value *X*_*i*_:

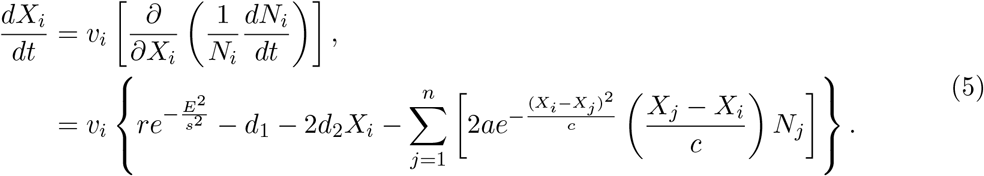

The rate of trait change is scaled by *v*_*i*_, the adaptation rate, which is assumed to be the only species-specific parameter within each community. This may also be termed the additive genetic variance, in the case that adaptation is genetic (Cortez and Patel 2017; Yamamichi and Letten 2021).

In a given environment, *E*, and in the absence of competition, there exists an optimal trait value, 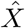, given by:

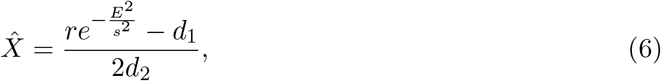

and a limited range of trait values that allow positive growth:

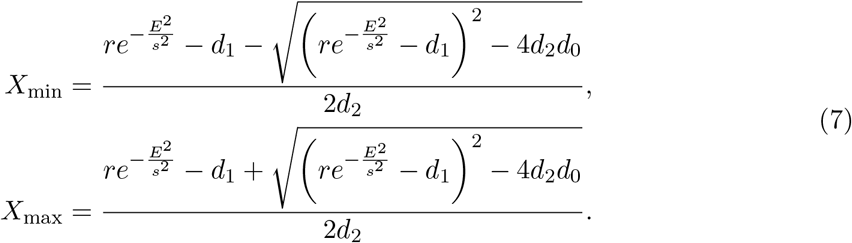

Note that, since *r, d*_1_, and *d*_2_ are not species-specific, 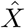 is the same for all species. The use of this quadratic function, given that *r > d*_1_ and *d*_2_ ≠ 0, implies that the trait confers a linear benefit to growth, while incurring a higher-order cost (Figure S1).

### Numerical experiment

We performed simulations of *in silico* competitive communities and observed how changes in the summed community population (functioning) and similarity to the pristine-condition community (composition) were impacted by environmental change, depending on the amount of environmental change (*E*) and the average adaptation rate of the community 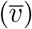.

We varied two factors when performing the simulations (Table 1).

**Table 1:** Factors varied in the simulations.

| Factor | Meaning | Levels |
| --- | --- | --- |
| $E$ | Amount of environmental change | 0.0625, 0.125, 0.25, 0.5 |
| $\bar{v}$ | Mean adaptation rate within community | 0, 0.01, 0.1, 1 |

For each of the factors varied (Table 1), we randomly generated 1,000 communities and simulated the community dynamics, totalling 16,000 simulations, in Julia 1.12.1 (Bezanson et al. 2017) using the DifferentialEquations package (Rackauckas and Nie 2017) with an implicit, stiff solver, Rodas5, (Di Marzo 1993) with relative and absolute tolerances of 10^*−*9^.

Each simulation was initialised with population abundances distributed normally around *K/n* of their carrying capacities, *N*_0_ *∼ K/n N* (1, 0.2), and the initial trait values distributed evenly around the unstressed trait optimum 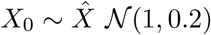.

Each community was first simulated until the it stabilised to some “assembled” state. During this assembly phase, the species densities, *N*_*i*_, settled into some equilibrium and species with *v*_*i*_ *≥* 0 underwent adaptation, with their traits, *X*_*i*_, changing to optimise growth. Following the assembly phase, we then altered the environment by changing *E* to one of the values in Table 1, and, keeping the state values from the final time point in the assembly phase, continued the simulations until they stabilised again.

### Parameterisation

All communities were initialised with eight species, *n* = 8, the interaction scaling factor *a* = 1, and the interaction variance *c* = 0.2. The remaining parameters were randomly generated each simulation.

The parameters responsible for mortality were randomly sampled from truncated normal distributions: 0.025 *< d*_0_ *∼ N* (0.075, 0.025), 0.1 *< d*_1_ *∼ N* (0.5, 0.1), and 0.025 *< d*_2_ *∼ N* (0.05, 0.05). For each community, *r* was computed numerically (using Newton’s method, NewtonRaphson, using the NonlinearSolve package in Julia (Pal et al. 2026) such that the carrying capacity of a species with the optimal trait value 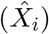 in a pristine environment (*E* = 0) was equal to 1 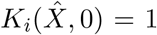, Equation 4). On average, this resulted in an *r* value of *≈* 1.07. The sensitivity, *s*, to environmental change, was randomly sampled from a normal distribution, 0.25 *< s ∼ N* (1, 0.1). The species adaptation rates, *v*_*i*_, were randomly distributed around 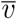 with a standard deviation of 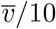 i.e. a coefficient of variation of 1*/*10.

Note that, within each community, only the *v*_*i*_ values were species-specific. All other parameters (*r, d*_0_, *d*_1_, *d*_2_, *s, a*, and *c*) were constant and any differences between species in their growth dynamics was due entirely to their trait values, *X*_*i*_.

### Simulation analyses

Each simulation produced a time series of species abundances and trait values, firstly in pristine conditions until *t*_change_ (assembly phase) and then at some level *E* until *t*_end_ (recovery phase). Using the final trait values, *X*_*i*_(*t* = *t*_end_), we firstly quantified the effect of the environment on the species intrinsic growth rate, *µ*(*E, X*_*i*_). We calculated both the mean response as 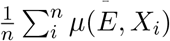 and the response variation as SD *µ*(*E, X*_*i*_). These two variables describe the effect of the environmental change on the community growth.

We used the full time series to quantify functional and compositional stability. Functioning was defined as the summed abundance of all species at each time point, and composition was defined using the Bray-Curtis similarity of the relative species abundances compared to that of the pristine assembled community at *t*_change_ ((Bray and Curtis 1957)Bray & Curtis 1957). We used relative abundances to avoid it trivially co-varying with functioning and, consequently, composition varied between 0 and 1.

We quantifed stability using the definitions of Van Meerbeek, Jucker, and Svenning 2021.

Resistance was quantified as:

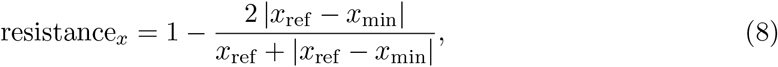

where *x* is either functioning or composition, *x*_ref_ is the value at *t* = *t*_change_ and *x*_min_ = min *x*_*t*_ *∀t > t*_change_. Resistance indicates how affected a variable is by the environmental change, relative to its initial value. Resistance of 1 indicates that the measurement is unaffected by the environmental change (*x*_min_ = *x*_ref_), while resistance of 0 indicates a 100% relative change of functioning/composition (*x*_min_ = 0).

We quantified resilience as:

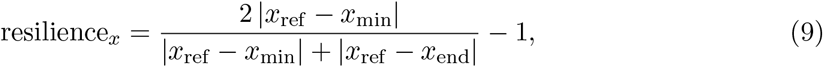

where *x*_ref_ and *x*_min_ are as before, and *x*_end_ is this value at the end of the simulations. Resilience of 1 indicates complete return to the assembled conditions by the end of the time series (*x*_end_ = *x*_ref_), while a resilience of 0 indicates that no change occured after reaching *x*_min_ (*x*_end_ = *x*_min_).

Finally, although similar in meaning to resilience, we also quantified recovery as:

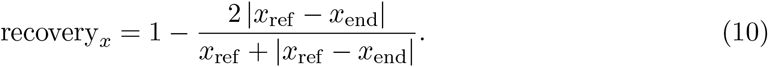

This is identical to the equation for resistance but quantified at the end of the simulation, indicating how close to pristine conditions the community returned, without any reference to the minimum reached.

Throughout the analyses, all confidence intervals were computed by bootstrapping the data 10,000 times.

## Results

### Effect of adaptation on species growth rates

A basic analysis of this eco-evolutionary community model showed that environmental change, *E*, reduces the maximum growth rate and shifts the optimal trait value downward (Figure 1 B, derived from Equation 6). This is because the growth rate declines quadratically as *X*_*i*_ moves further away from 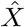 and changing the environment *E* causes a reduction in carrying capacity, a reduction of the optimum trait value, and a narrowing of the viable trait range (Figure 1 A). Since the effect of trait on growth is quadratic and the rate of trait change depends on the derivative of the growth rate with respect to *X*_*i*_, the rate of trait change is linear with respect to *X*_*i*_ (Figure 1 B). A consequence of this result is that increasing *E* compresses the viable trait range and lowers the trait optimum (Figure 2).

**Figure 1:**
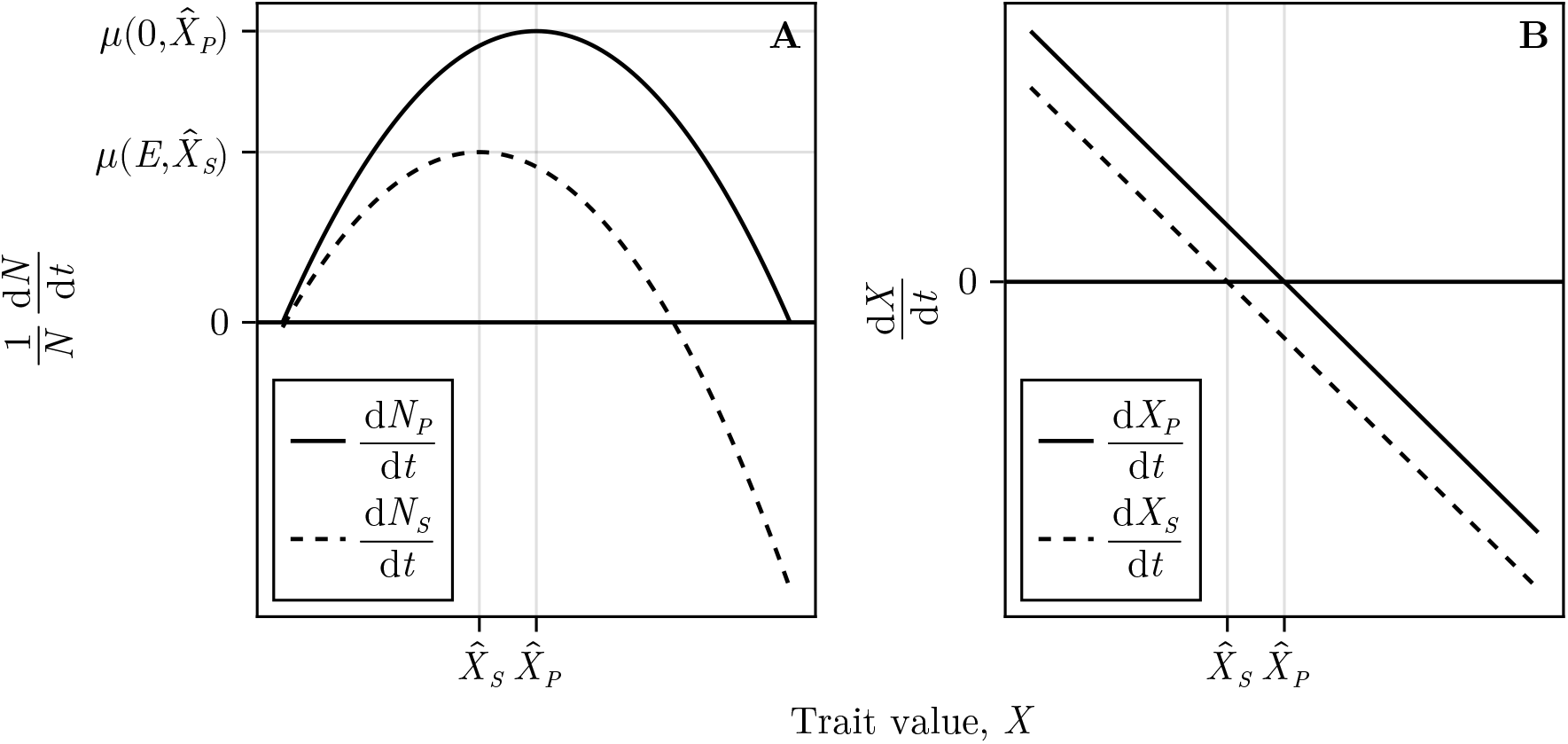
The effect of the trait value, *X*, on 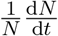 and 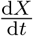 in both pristine (solid line) and stressed (dashed line) environments i.e. *E* = 0 and *E ≠* 0, respectively. The maximum population growth rate in pristine conditions, 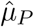 , is higher than in stressed conditions, 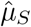 and the pristine trait optimum, 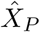 , is higher than the stressed trait optimum, 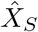. Parameters used: *r* = 1.415, *d*_0_ = 0.0732, *d*_1_ = 0.553, *d*_2_ = 0.173, *a* = 1, *c* = 0.2, and *s* = 1.116.

**Figure 2:**
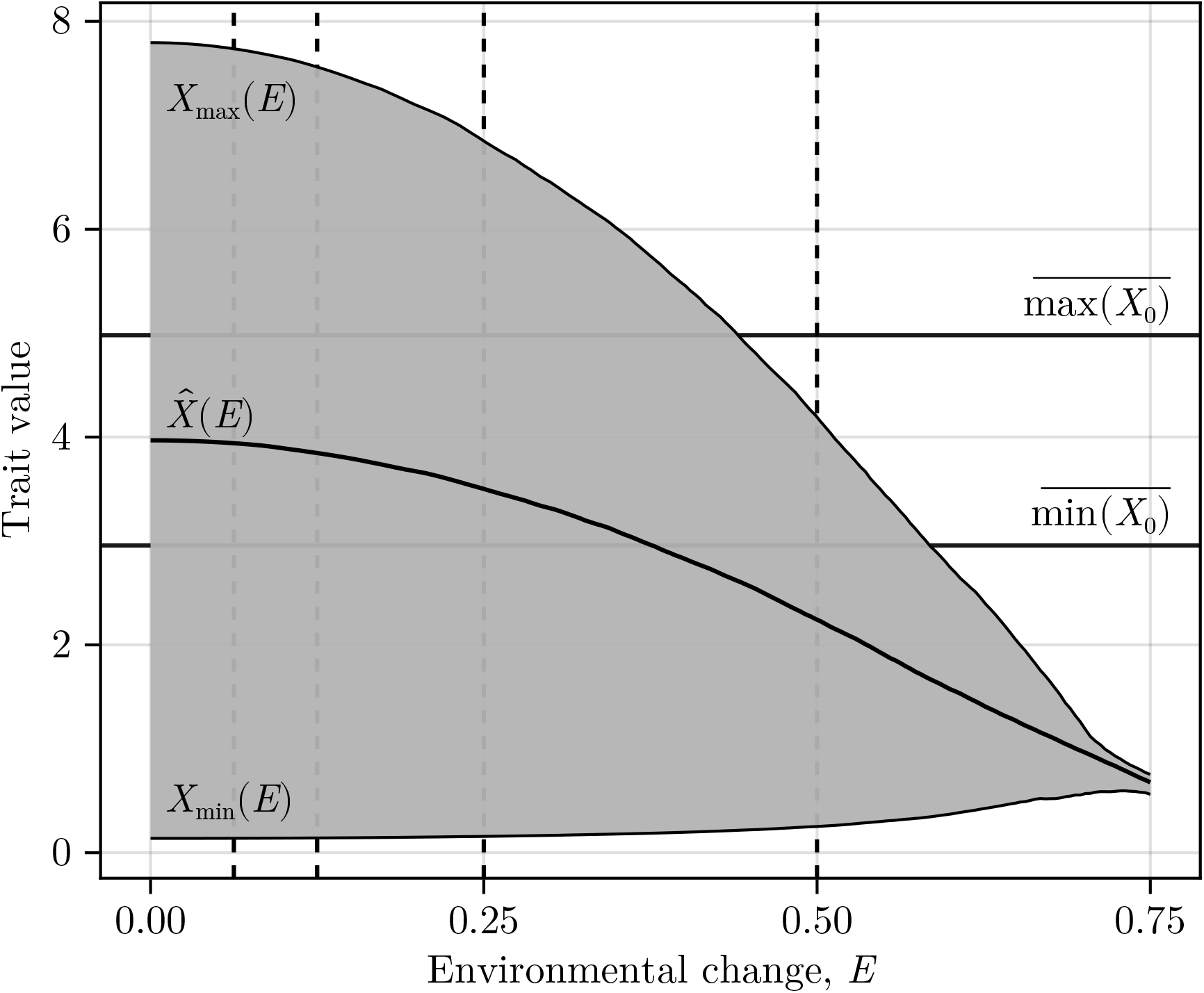
The range of viable trait values across a gradient of environmental change, *E*, derived from Equation 7. The solid black line indicates the trait optimum 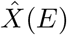 and the grey region is the viable range of trait values between *X*_min_(*E*) and *X*_max_(*E*). The vertical dashed lines indicate *E* = 0.0625, 0.125, 0.25, and 0.5 (the values used in the simulations). The horizontal dashed lines indicate the average minimum and maximum starting trait values, *X*_0_ = *X*(*t*), *t* = 0, used in the simulations. Parameters used are the same as in Figure 1.

The simulations showed that, at low *E*, traits in non-adaptive communities remained clustered near the pristine optimum, giving high average growth but low variation (Figure 3). As *E* increased, species further from the stressed optimum had lower growth rates, increasing this variation. In adaptive communities, by contrast, the compression of the viable trait range shaped growth rates differently depending on *E*: at low *E*, traits diverged to reduce competition, lowering average growth but increasing its variation; at high *E*, the shrinking viable range forced traits back together, reducing variation. At the high end of environmental change levels considered here, *E* = 0.5, adaptive and non-adaptive communities showed similar levels of response variation.

**Figure 3:**
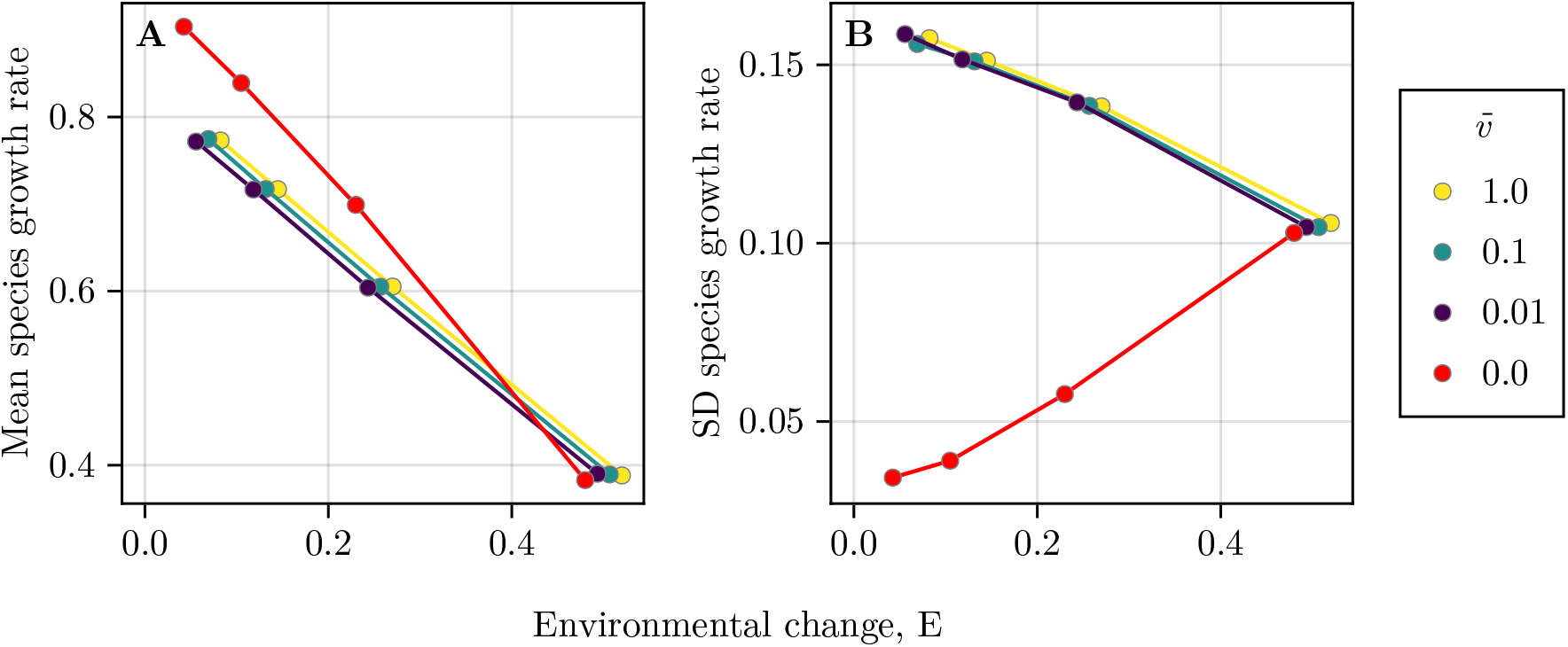
The effect of environmental change (x-axis) and community adaptation rate (colour) on mean species growth rate, 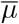, (A) and the standard deviation of growth rates, SD(*µ*), (B). Points indicate mean values and error bars indicate bootstrapped 95% confidence intervals, across 1,000 simulations.

### Implications for stability: Illustration

We firstly present two example communities, with and without adaptation, to demonstrate a typical simulation trajectory (Figure 4). Following the onset of environmental change, species abundances in the non-adaptive community rapidly settle to a new equilibrium while this process is slower in the adaptive community (Figure 4 A and B). The traits in the non-adaptive community remain constant while they slowly settle at lower values in the adaptive community (Figure 4 C and D). These dynamics have consequences for total abundance (i.e. the function considered; Figure 4 E and F) and community composition (i.e. Bray-Curtis similarity to the assembled community; Figure 4 G and H). Specifically, functioning and composition either quickly settle (without adaptation), or show transient behaviour that involve sharp drops relative to the pre-change value (with adaptation). These results, in turn, have consequences for stability calculations, which rely on the values for function and composition at specific timepoints (black points; Equation 8, Equation 9, and Equation 10).

**Figure 4:**
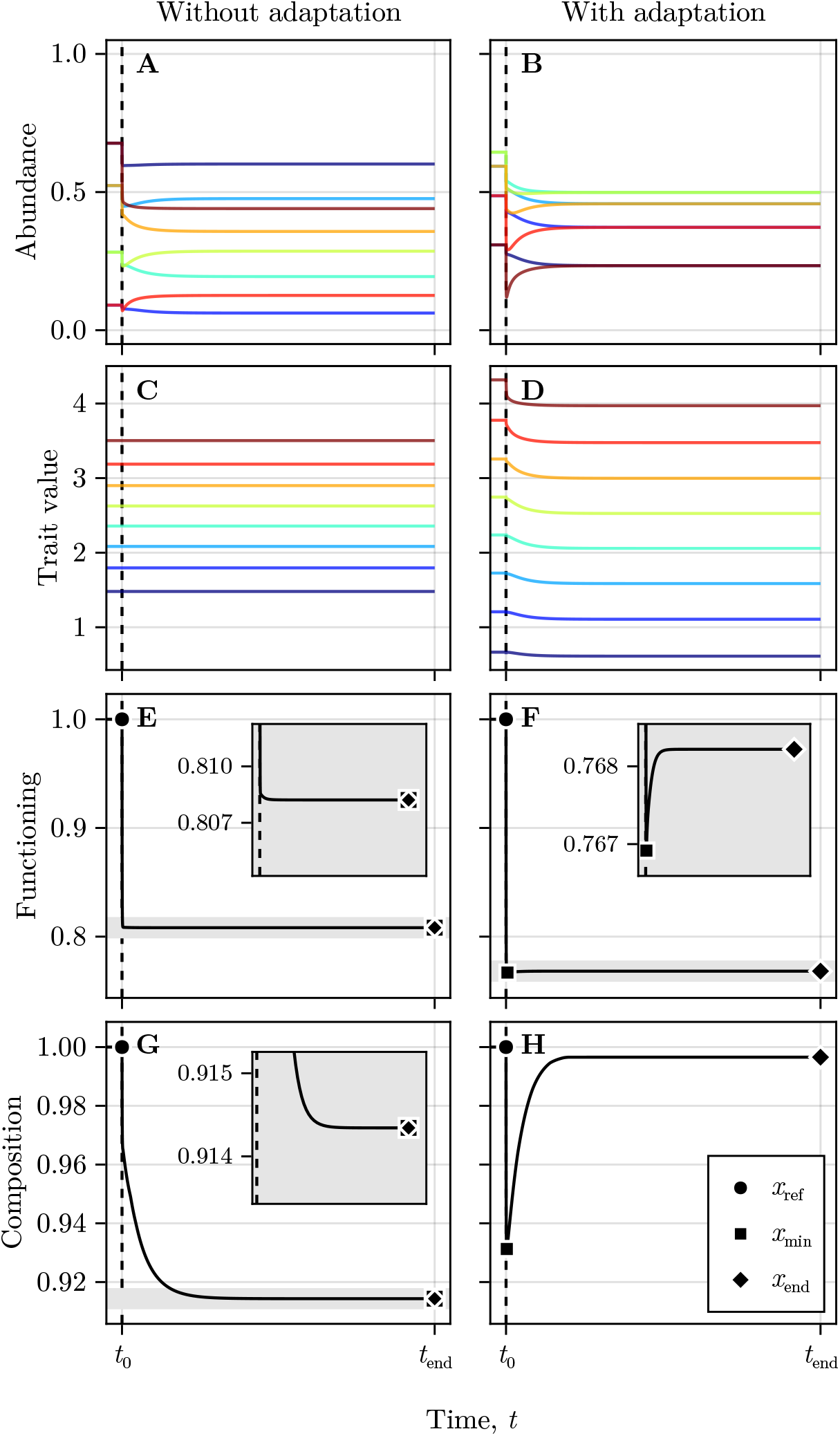
Demonstration of community abundances (A and B), trait values (C and D), and functioning (E and F), and composition (G and H) for a demonstration community without (left column) and with (right column) adaptation. The onset of environmental change is indicated by the vertical dashed line. Different species are indicated by colours. Parameters used are the same as in Figure 1.

Figure 5 illustrates these mechanisms in the trait-abundance space, and shows the importance of adaptation rate, 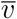. Following environmental change, species abundances declined from their assembled state, while traits adapted towards the new stressed optimum, 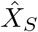 , eventually settling into a new community structure. For species to the right of the new trait optimum, this drop in abundance was greater when adaptation was slower, which increased the transient return toward the pre-disturbance state, and therefore increased the measured resilience metric, both of community composition and abundance. Similar results have been observed in studies of evolutionary rescue, which can result in similar species recovery trajectories (Orr and Unckless 2014). Recovery was largely independent of adaptation rate, because 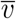 affected the speed of trait change but not the post-stress equilibrium trait values. The less-adaptive populations did, however, fall to lower abundances (i.e. lower resistance) prior to recovering to the same final state (i.e. higher resilience but equal recovery). In non-adaptive communities, traits remained fixed and species dominance changed little, so functional and compositional resilience and recovery were negligible or absent.

**Figure 5:**
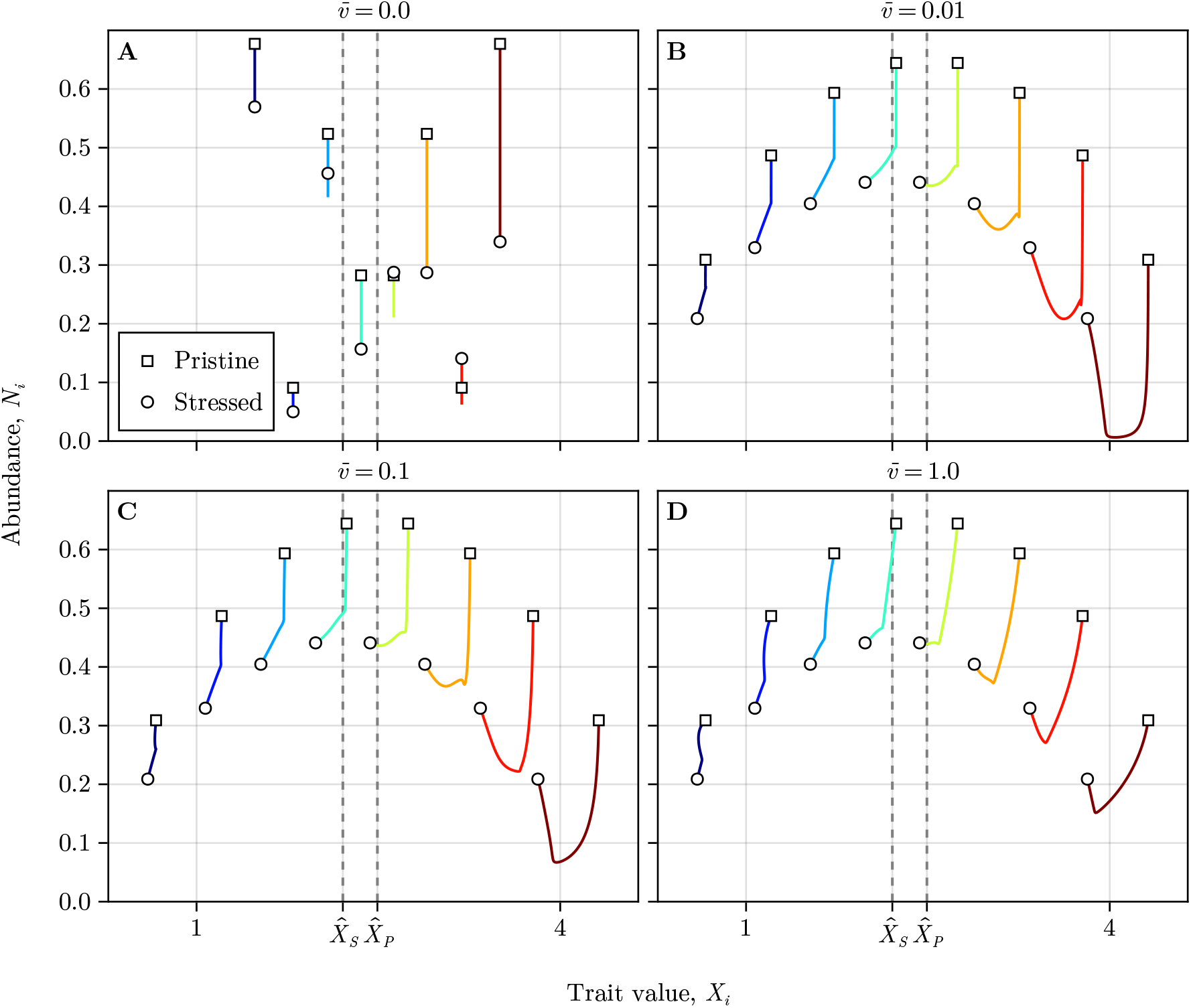
Demonstration of community dynamics for different values of 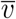 (panels). The vertical dashed lines indicate the unstressed and stressed optimum trait values. Different species are indicated by colours. The squares and circles indicate the assembled community and the post-stress adapted community, respectively. Apart from 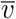, the parameters and initial conditions are the same as in Figure 4.

### Implications for stability: Simulations

The illustrated consequences of trait adaptation for the three stability metrics were prevalent in our simulation results (Figure 6). Adaptation led to slightly reduced functional resistance but improved compositional resistance.

**Figure 6:**
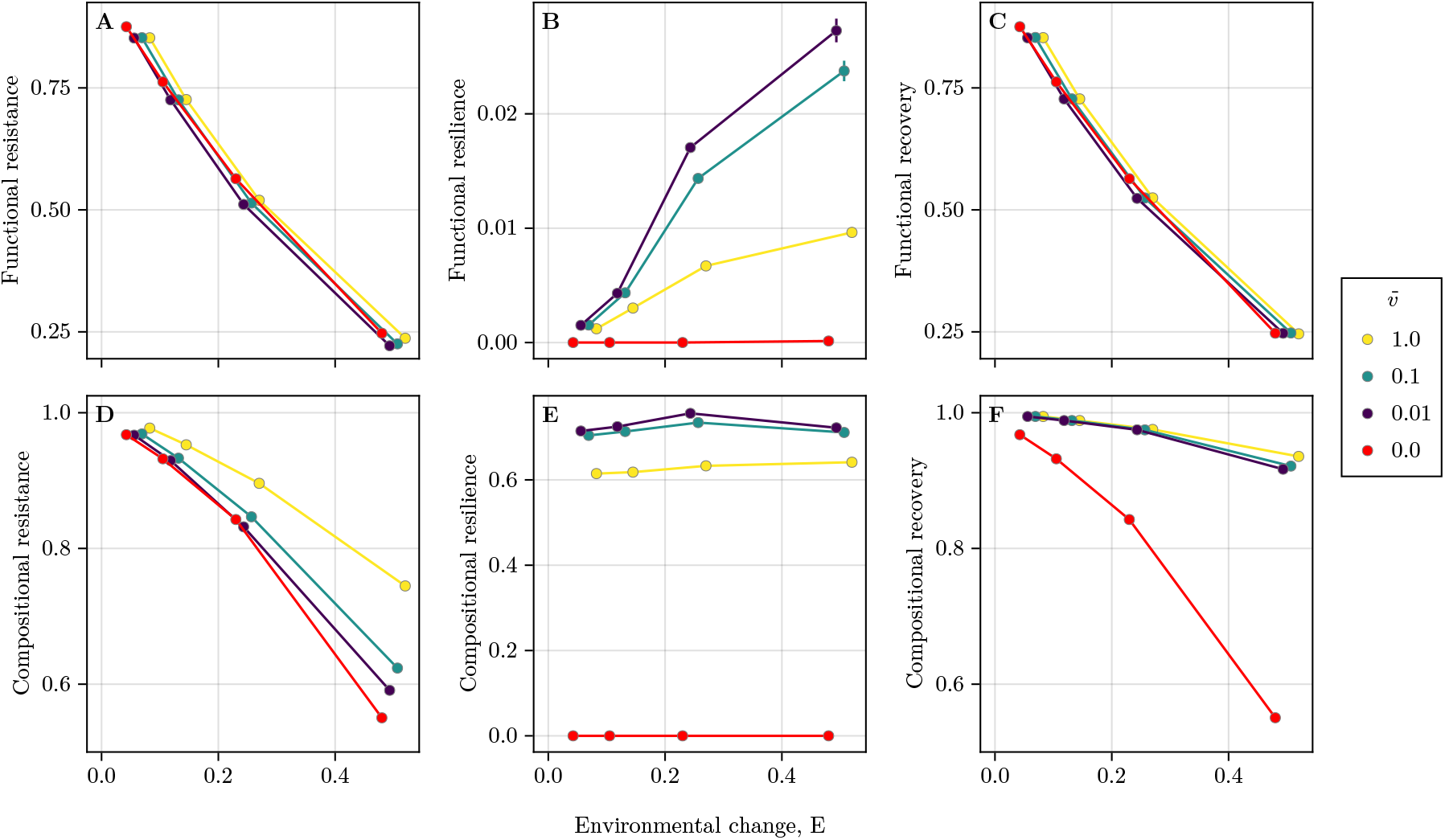
The effect of environmental change (x-axis) and community adaptation rate (colour) on the 6 stability metrics considered (y-axes): functional resistance (A), resilience (B), recovery (C), compositional resistance (D), resilience (E), and recovery (F). Points indicate mean values and error bars indicate bootstrapped 95% confidence intervals. Note that many of the error bars are too small to visible.

Trait adaptation had a slightly negative effect on functional resistance for most levels of environmental change but strongly improved compositional resistance (Figure 6 A and D). Any amount of adaptation enabled resilience and recovery under sustained environmental change (Figure 6 B, C, E, F), while non-adaptive communities showed no resilience of either functioning or composition. Adaptive communities were much more compositionally-than functionally-resilient and, notably, resilience was lower at higher rates of adaptation (Figure 6 B, E). The recovery of adaptive communities, however, was more similar across rates of adaptation (Figure 6 C and F). That is to say that adaptive communities recovered to approximately the same point eventually, but that less-adaptive communities fell further during this process. There was also a slightly positive effect of 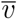 on compositional recovery - lower resilience was offset by greater recovery.

## Discussion

Differences between non-adaptive and adaptive communities in mean response and response variation (Figure 3) reflected the balance between environmental optimisation and minimizing competition. In near-pristine conditions, where *E ≈* 0, competition was a stronger selective force than environmental mismatch. In adaptive communities, this favoured trait divergence, which reduced overlap in competitive effects but also moved some species away from the environmental optimum, thereby increasing variation in species growth responses. By contrast, as the non-adaptive communities were initialised relatively close to the trait optimum, 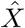, remained well-matched to the unchanged environment, and therefore showed higher mean intrinsic growth and lower response variation. Under stronger environmental change, however, the viable trait range narrowed, forcing species closer together and constraining the extent to which competition could drive divergence. As a result, response variation declined in adaptive communities as *E* increased. Taken together, these results show that competition was a major determinant of how adaptation shaped community responses to environmental change. This is consistent with theoretical work showing that competition and trait evolution jointly determine community-level responses to environmental gradients (Barabás et al. 2022; Barabás and D’Andrea 2016), and with empirical evidence that competitive interactions can modulate species responses to environmental change (Van Dyke, Levine, and Kraft 2022).

Our results show that trait adaptation altered functional and compositional stability asymmetrically (Figure 6). Adaptation consistently promoted compositional stability, whereas its effects on functioning were weaker and depended on the stability component considered. In our model, the magnitude of environmental change was the main determinant of functional loss, and increasing adaptation rate only modestly altered functional resistance. By contrast, compositional stability responded much more strongly to adaptation. This pattern is consistent with earlier results that functional change depends on the mean response across species, whereas compositional change depends on variation in responses among species (Holmes, Spaak, and De Laender 2021). Thus, adaptation did not overturn this distinction, but instead acted mainly by reshaping the distribution of species responses. More broadly, this asymmetry is consistent with theoretical and experimental work showing that eco-evolutionary dynamics can reshape community composition more readily than ecosystem-level productivity (Ellner, Geber, and Hairston 2011; Hairston et al. 2005; Turcotte, Reznick, and Hare 2011).

At high levels of environmental change (*E* = 0.5) the adaptive and non-adaptive communities experienced the same response mean and variation (Figure 3), however, the adaptive communities were still substantially more compositionally stable in all metrics considered (Figure 6). Despite species being differently affected, the species traits adapted in unison to maintain the original community trait structure, just shifted downwards and less widely-dispersed. The same response mean and variation can result in drastically different consequences depending how these species are organised in the trait space.

Increasing the rate of adaptation conferred greater resistance but lower resilience. Faster-adapting communities experienced smaller immediate changes in abundance and composition following environmental change, and were therefore more resistant. At the same time, because they were displaced less from their pre-disturbance state, they showed less subsequent return towards that state, and therefore lower resilience under our metric. Slower-adapting communities showed the opposite pattern. The faster the community adapts towards the new optimum, the smaller the transient decline in abundance, and hence the smaller the subsequent “rebound” that resilience can capture (Figure S2). Importantly, this does not mean that faster adaptation made communities qualitatively less stable. Rather, because, resilience is measured relative to the initial displacement, reducing the initial impact of environmental change necessarily leaves less return to quantify. Resistance and resilience therefore did not simply co-vary as interchangeable expressions of stability, but responded differently to adaptation rate. This adds to growing recognition that different stability components can respond in contrasting ways to the same ecological or evolutionary process (Baert et al. 2016; Radchuk et al. 2019), and suggests that empirical assessments should be cautious in interpreting resilience without considering the magnitude of the initial disturbance response (Donohue et al. 2016; Van Meerbeek, Jucker, and Svenning 2021).

The difference between resistance and resilience is crucial in communities that are subject to stochasticity. The more time spent at low abundance, the greater the likelihood of stochastic extinction (Catullo et al. 2019). With lower rates of adaptation, slow transient states that leave some species vulnerable to extinction are more likely, quantifiable using an integrated stability metric (Hastings et al. 2018; Urrutia-Cordero et al. 2022).

Adaptive communities displayed resilience even though the environmental change was maintained throughout the simulation. Because the stressor did not relax, any return towards the pre-disturbance state could not arise from restoration of the original external conditions. In non-adaptive communities, traits remained fixed and abundances settled to a new equilibrium displaced from the pre-disturbance state, so resilience and recovery were negligible or absent. Adaptation altered this outcome by shifting traits towards the new optimum, which increased species growth rates and allowed partial recovery of lost abundance and composition. The resulting dynamics differ from classical engineering resilience (Pimm 1984), which typically describes return following relaxation of a perturbation. Instead, the resilience observed here arose because species adjusted their internal state in response to the changed conditions. In that sense, the mechanism is closer to adaptive capacity (Gunderson 2000; Van Meerbeek, Jucker, and Svenning 2021) or, at the population level, to evolutionary rescue (Bell and Gonzalez 2009; Gomulkiewicz and Holt 1995), here expressed at the community scale. One implication is that empirical measurements of resilience in systems capable of rapid trait change may partly reflect adaptive recovery rather than ecological return dynamics alone, and the two mechanisms may be difficult to disentangle from time-series data.

These findings contribute to a growing body of work showing that rapid evolution, occurring on timescales comparable to ecological dynamics, can have measurable consequences for community and ecosystem properties (Ellner, Geber, and Hairston 2011; Hairston et al. 2005). While other studies have focused mainly on single-species systems or simple pairwise interactions (Yoshida et al. 2003), it is now clear that eco-evolutionary feedbacks can shape multispecies dynamics in ways that cannot be predicted from ecological processes alone (Yamamichi, Gibbs, and Levine 2022). Our results extend this perspective by showing that the rate of adaptation, not only its presence or absence, influences which stability components are most strongly expressed. More specifically, in our model, competition-driven trait divergence during assembly likely increased niche differences among species, helping maintain diversity in the assembled community and shaping how that community subsequently responded to environmental change. This is consistent with theoretical and empirical work showing that character displacement can stabilise coexistence by reducing competitive overlap (Barabás and D’Andrea 2016). It also suggests that some of the compositional stability benefits of adaptation observed here depend on the eco-evolutionary history of community assembly. Communities assembled under weaker competitive sorting, or with less scope for rapid trait change, may therefore show weaker compositional stability under environmental change.

In ecotoxicology, conservation biology, and climate-impact assessment, communities are often modelled with fixed species responses, ignoring adaptation (Mouquet et al. 2015). Our model shows that omitting adaptation can lead to a slight overestimation of functional resistance but underestimation of compositional resistance and of recovery, especially compositional recovery. Non-adaptive models would therefore tend to portray communities as compositionally more fragile than they may be when rapid trait change is possible, while only slightly misrepresenting the immediate decline in aggregate functioning. This pinpoints a broader problem in applied ecology: predictive approaches frequently omit evolutionary responses to environmental change and can therefore misestimate persistence, range shifts, or community reassembly (Cotto et al. 2017; Urban et al. 2016).

A potential extension of the approach presented would to explore additional regimes of environmental change, such as gradual change or across multiple pulses, rather than as a single discrete event. Additionally, our model assumes all species share the same trait-growth relationship and differ only in their adaptation rate and initial trait value. Relaxing this assumption by, for instance, allowing species-specific environmental sensitivities or trait optima, would allow the model to represent the full range of response diversity documented empirically (Ross et al. 2023), and may reveal further interactions between response diversity and adaptation rate in shaping stability.

The effects of rapid adaptation on community stability, even in this highly-simplified model, were not immediately intuitive. The contrasting needs to reduce competition and optimise growth in stressed environment caused response variation to co-vary with environmental change in adaptive comminities — the inverse of the pattern observed in non-adaptive communities. Higher rates of adaptation shifted communities from resilient to resistant, meaning that in addition to a reduced total impact of environmental change on functioning, species should also be less vulnerable to stochastic extinction. Furthermore, the ability to adapt at a rate comparable to ecological processes meant that communities were more compositionally stable overall, with species shifting their traits in unison. Adaptation, however, is not capable of completely mitigating the effects of environmental change entirely, and abiotic constraints on growth can override the benefits of adaptation in highly altered environments.

