## Supplementary figures for "Rapid trait adaptation has contrasting effects on different dimensions of stability under environmental change"

### Supplementary materials

October 2, 2026

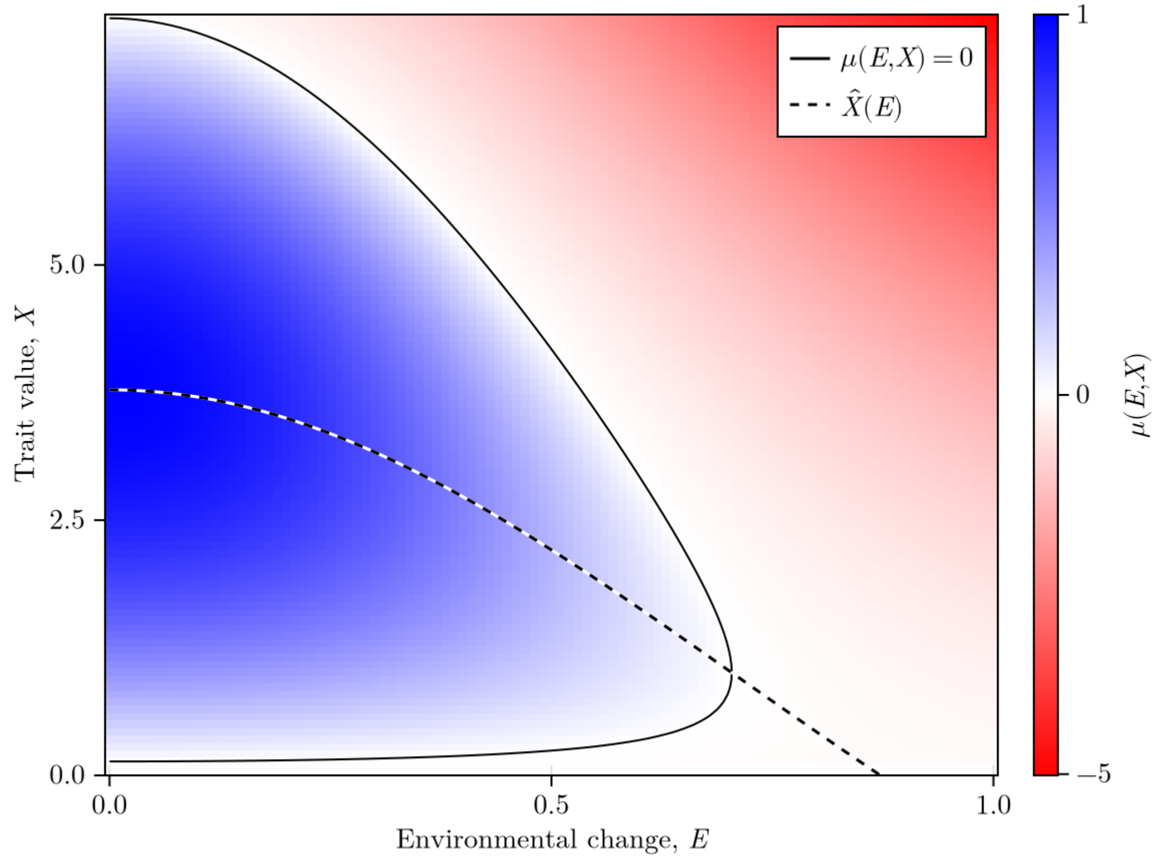


Figure S1: The effects of $E$ and $X$ on growth.

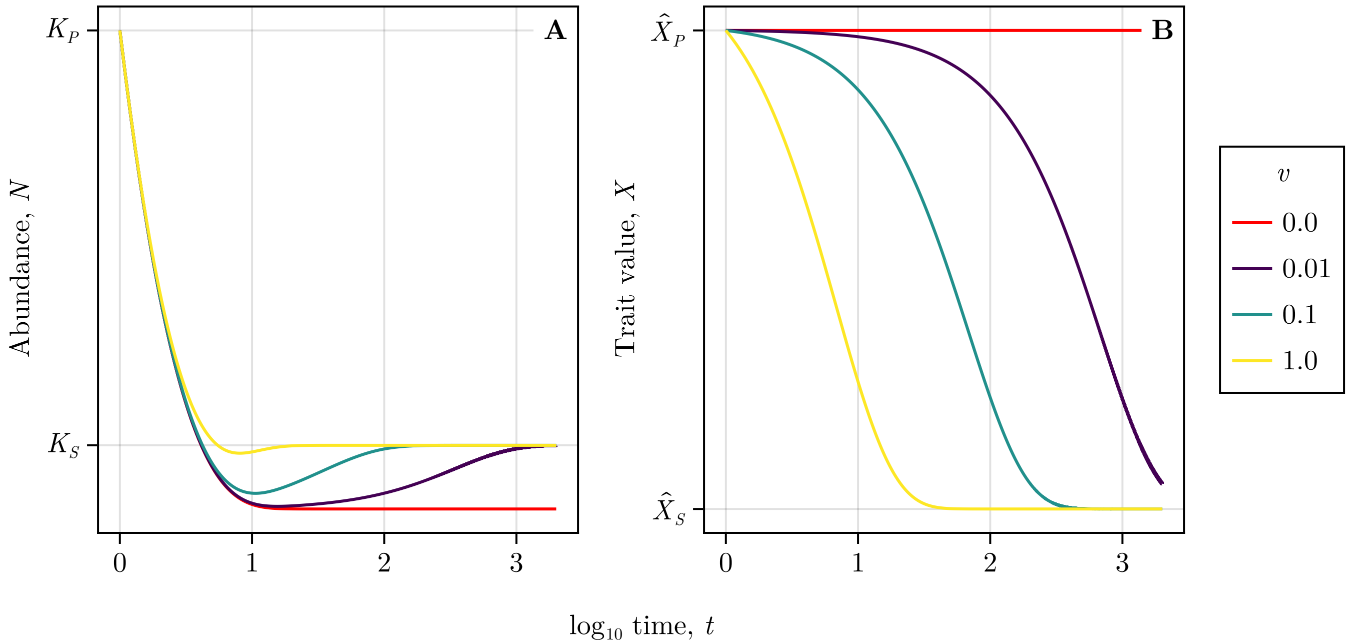


Figure S2: The effect of adaptation rate on abundance (A) and trait value (B)
